# A Visual, Inexpensive, and Wireless Capillary Rheometer

**DOI:** 10.1101/2022.09.25.509390

**Authors:** Jianyi Du, Soham Sinha, Stacey Lee, Mark A. Skylar-Scott

**Author notes:** **Corresponding author’s email address and Twitter handle**, Twitter: @mascott85.

## Abstract

Complex fluid systems can exhibit highly non-linear and time-dependent flow behaviors that arise from microscale reorganization. Rheological measurements to understand these flow-induced properties in soft condensed matter and biology are critical to optimize manufacturing processes that rely upon the use of complex fluids, such as the creation of viscoelastic bioinks for 3D bioprinting. While typical rheological characterizations are performed on bulky and expensive rheometers, these options do not readily enable *in situ* microscopic visualization of flow behavior, which can provide microstructural insights into the mechanisms and patterns of yielding. To address these limitations, we present a visual, inexpensive (approximately $200), and wireless capillary rheometer (VIEWR) assembled from chiefly 3D printed components. We validate the accuracy and reliability of the portable rheometer by comparing measurements of multiple rheological parameters, including viscosity and yield stress, with measurements obtained from a commercial oscillatory rheometer. At its core, VIEWR employs an easily interchangeable glass capillary channel to enable real-time microscopic observation under flow conditions. To facilitate various experimental conditions, a communication protocol based on Internet of Things (IoT) is adapted to support wireless transmission of data. This unique rheometer design can be used for characterizing and visualizing numerous living and non-living complex fluids.

**Specifications table:** 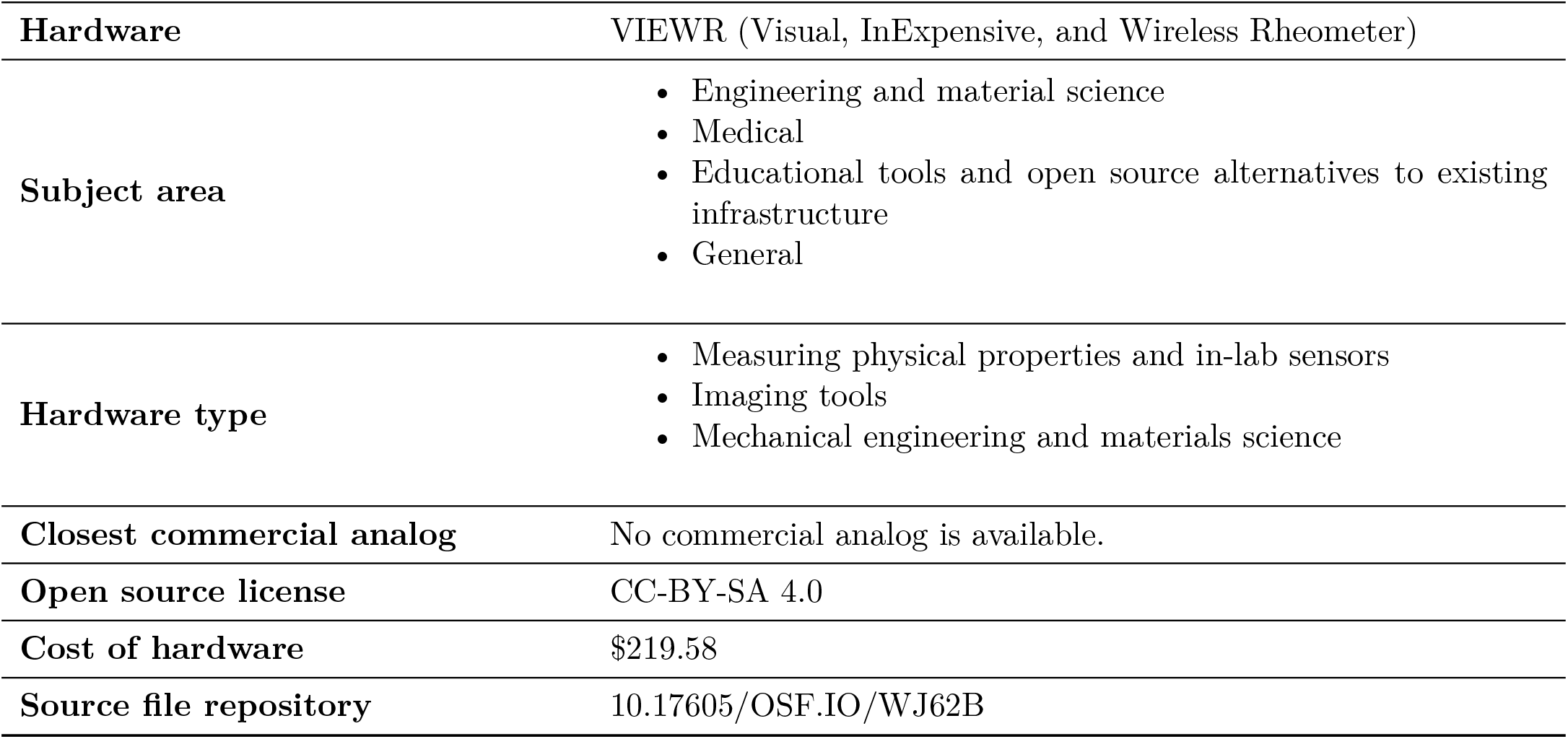

## 1. Hardware in context

Rheology is the study of how materials flow under applied stress. By applying and measuring forces and deformations, a rheometer can extract key material properties of fluids such as their viscous and elastic behaviors under small or large deformations [1]. Rheometers are used by scientists, engineers, and technicians across many different industries, and are found in tens of thousands of laboratories that study a wide range of disciplines including polymer science [2, 3], biosciences [4], manufacturing [5, 6], food [7], energy [8], and cosmetics [9].

There are two major types of commercial rheometers: rotational (or oscillatory) rheometers, and capillary rheometers. Rotational rheometers measure fluid properties by sandwiching samples between two co-axial surfaces, one stationary and one rotating, thereby inducing a well-defined shear deformation while measuring the applied force or torque [1]. Conversely, a capillary rheometer flows samples through a narrow capillary tube driven by a defined shear rate or pressure, and the fluid properties can be quantified by measuring the pressure gradient or flow kinetics [10] (Fig. 1a). While rotational rheometers can perform more comprehensive sets of measurements, capillary rheometers are far simpler and cheaper to construct, operate, and maintain, while being able to extract many important material properties of different complex fluids. By applying a flow ramp, a fluid’s viscosity, or its resistance to external flow, can be measured to identify whether the material exhibits shear thinning (such as blood [11] or paint [12]), shear thickening (such as cornstarch suspensions, also known as Oobleck [13]), or shear-independent (such as Newtonian fluids, including water and honey) behavior. For a specific class of fluids, such as mayonnaise or ketchup, this flow ramp can be used to extract a shear yield stress, wherein the material does not flow until the applied shear stress exceeds this critical value [14]. Upon the removal of external shear stresses, many yield stress fluids “self-heal” or re-solidify over time due to the reconstruction of internal structure [15]. Typically, the kinetics at which materials yield or self-heal can be extracted by step-and-hold experiments wherein flow is repeatedly initiated and ceased (Fig. 1b).

**Figure 1:**
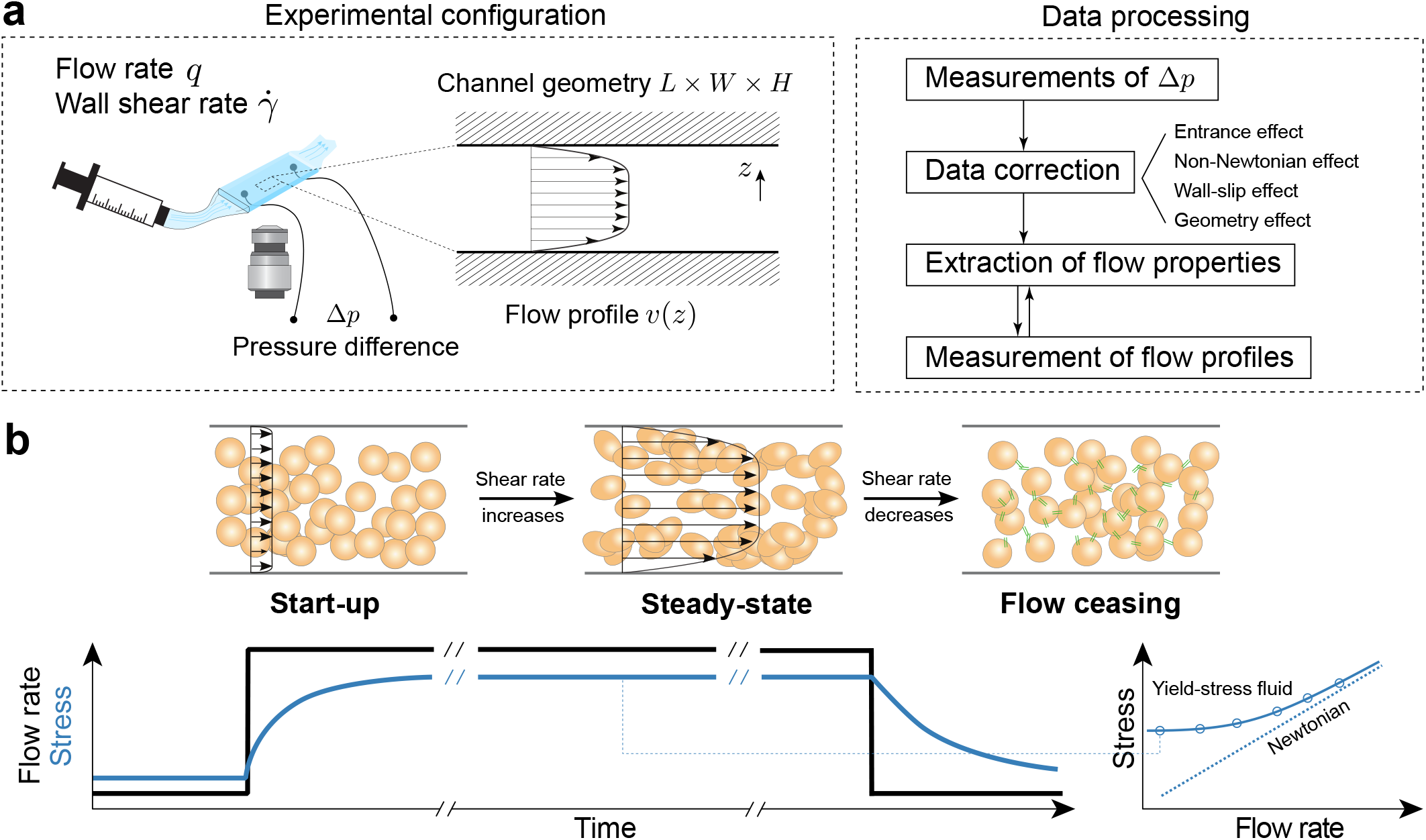
Overview of a capillary rheometer and its assays. (a) Schematic of capillary rheometry (with *in situ* observation and flow velocimetry) and data processing. (b) Typical flow measurements performed on a capillary rheometer. Right: Flow curves of a Newtonian fluid and a yield-stress fluid.

In addition to their simplicity, capillary rheometers exhibit several advantages over the rotational counterparts in certain scenarios. With selected channel geometries, a capillary rheometer can measure fluid properties accurately at much higher shear rates than a commercially available rotational rheometer (which is generally capped at 500s^−1^ to 1000s^−1^ due to fluid inertia, secondary flow and motor speed limit [16]). In addition, for applications that require forced extrusion through narrow channels, such as 3D printing, a capillary rheometer can measure and optimize ink-flow behaviors in a manner that more closely resembles its end-use. Finally, while existing commercial capillary rheometers do not incorporate optical components, the use of a transparent capillary channel can facilitate real-time microscopy for rheo-optic measurements of flow-induced properties [17, 18]. Our laboratory is principally interested in understanding the forces and flows associated with extrusion 3D printing. In a typical extrusion printing process, the ink flow in the printing nozzle is initiated by a sufficiently large pressure difference. This specific flow type, identical to the flow in a capillary rheometer, can be principally described by a Poiseuille flow. In a simplified two-dimensional scenario, the dynamics of a Newtonian fluid are governed by the Hagen-Poiseuille equation [1] as

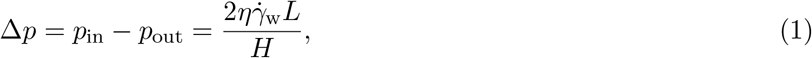

where *η* and 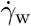 are the shear viscosity and the imposed wall shear rates, and *L* and *H* are the distance between the two pressure sensors and the height of the capillary channel, respectively. For a Newtonian fluid, the wall shear rate can be connected to the imposed flow rate per unit width via 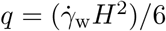 [1]. In practice, accurate measurements of the material functions are obtained through Eq. 1 with necessary mathematical corrections in the post-processing to account for the complex non-Newtonian material behaviors and experimental conditions that deviate from the hypotheses of a 2D simplification [19] (Fig. 1a).

In addition to rheological measurements at larger length scales, capillary rheometry can be readily combined with high-resolution imaging to study the variation and patterns of complex ink yielding under certain flow conditions. Particularly, in an extrusion-based bioprinting process, cell-encapsulated bioinks undergo non-linear deformation via shear and extensional flow through a nozzle geometry to form a desired printed filament. Bioinks commonly feature non-trivial yield stresses and shear-thinning properties to ensure both printability and self-supported structures after printing [20]. The magnitude of the yield stress is key to printed shape fidelity, and can be primarily contributed from the characteristics of intra- and intercellular structures, which can be readily inferred from direct observation of the morphological evolution under an external strain or stress [21]. Furthermore, yield-stress materials exhibit plug flow within channels [22] (*i.e*., a solid core of material under yield stress does not shear within the nozzle). Within this non-yielding section, cells can be protected from necrosis due to large deformation [23]. An enhanced understanding of the morphology and kinetics of polymer inks and living bioinks that are subject to flow conditions during the printing process can assist in the design and optimization of an accurate and efficient bioprinting process for fabricating scalable tissue constructs to recapitulate the physiological features of native counterparts.

Current commercially available capillary rheometers, while working on an identical measuring principles, can be broadly categorized into metal-based systems (main brands include Instron^®^, Rosand^®^, Rheograph^®^, etc) and microfluidic-based systems (RheoSense^®^). However, these commercial options are often prohibitively expensive (with typical cost over $20,000). Metal-based instruments do not generally support *in situ* flow visualization, while microfluidic-based capillary rheometers that allow for microscopic observation are largely available only through in-house fabrication or customization [24]. In addition, the capillary channels on available instruments are non-disposable, and are subject to constant clogging and cross-contamination, which inhibits accurate measurements of rheologically complex systems. Furthermore, commercial instruments are commonly large in size and rely on cable-based communications, impeding their operation inside a sterile or environmentally controlled enclosure such as a biosafety cabinet or CO2 incubators, as are commonly used for maintaining living systems [25].

An ideal capillary rheometer should be low cost, feature single use channels, be compact and wirelessly controlled for sterile operation. Here, we present a Visual, Inexpensive, and Wireless Capillary Rheometer (VIEWR). This portable rheometer features 3D-printed components that can be integrated with an inverted microscope, and flow channels assembled from commercially-available glass capillary tubes that can be easily cleaned and replaced, allowing for accurate measurements and real-time observation under given flow conditions. In addition, we designed web-based graphical user interfaces (GUI) based on Internet of Things (IoT) techniques, which enables the monitoring and characterization of cellular materials under controlled conditions (temperature, humidity, sterility, etc) through wireless communications. Our prototype was validated using a selection of biomaterial systems. We visualized the flow profiles of the tested systems in the channel, and compared with the predictions based on independent rheological characterizations from a commercial rheometer. We show that this system can be readily applied to hydrogel and polymer systems that are regularly employed in the biofabrication field.

## 2. Hardware description

### 2.1 Overview

The VIEWR is composed of a 3D-printed framework, which houses a custom syringe pump as an extruder, an interchangeable rectangular glass capillary as the flow channel, two pressure sensors (upstream and downstream) to measure material properties, and necessary electronic components, including a stepper motor driver, and a microcontroller with battery-power supply (Fig. 2a). The liquid sample is loaded into a gastight glass syringe (Part no. 81460, Hamilton Company; other syringes can be used with the holder geometries modified accordingly), which is installed on a linear stage and fed into the inlet tower. The stepper motor translates the linear stage and imposes flow through the glass capillary and the gauge pressures at both inlet and outlet towers are measured and transmitted to the processor for the calculation of material properties.

**Figure 2:**
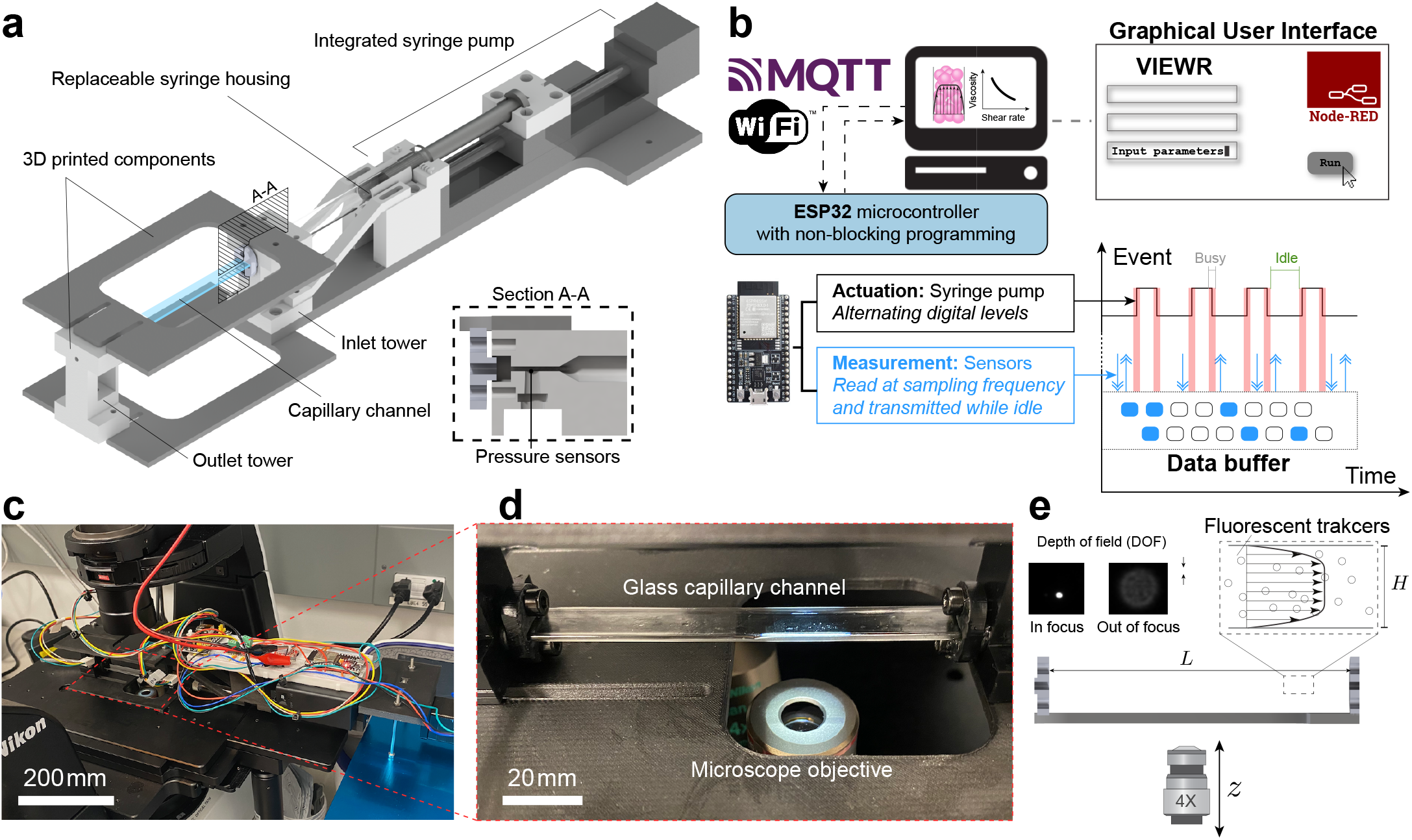
Design overview of the Visual, Inexpensive, and Wireless Capillary Rheometer (VIEWR). (a) Rendered assembly of the 3D-printed framework and key components, with the inset showing a section view of the inlet channel. (b) Instrument control scheme showing GUI, microcontroller, and the non-blocking method for interlacing actuation and reading. (c) Experimental configuration with the VIEWR installed on an inverted Nikon Ts-2R epifluorescence microscope. (d) Close-up view of the capillary channel installed over a 4X phase-contrast objective. (e) Schematic showing how an inverted microscope enables visualization of the flow profile in the capillary channel.

The 3D-printed structure is composed of a framework made of nylon reinforced with chopped carbon fibers (Markforged brand name Onyx™, dark gray), and liquid-handling components (inlet and outlet towers and syringe holders) made of stereolithography (SLA)-printed resin (B9R-2-Black, white gray). The rheometer can be installed onto an inverted microscope via two 3D printed adapters located on the top and bottom of the VIEWR (Fig. 2a). The microscope adapters also serve to provide additional support for the instrument. All components are assembled using hex screws and nuts. A commercially-available glass capillary (Catalog no. 63832-10, Electron Microscopy Sciences) connects the inlet and outlet towers to create a flow channel with a well-defined geometry and facile microscopic observation (Fig. 2e). To ensure shatter-free installation of the glass capillary, a custom-designed O-ring-enclosed connector is applied to adapt the glass capillary to the resin towers (inset of Fig. 2a, and Fig. 3b). The pressure difference between the inlet and outlet towers is measured by two analog gauge pressure sensors (model no. BPS130-HG015P-3S, Bourns Inc.) inserted from a sensor opening in each tower.

**Figure 3:**
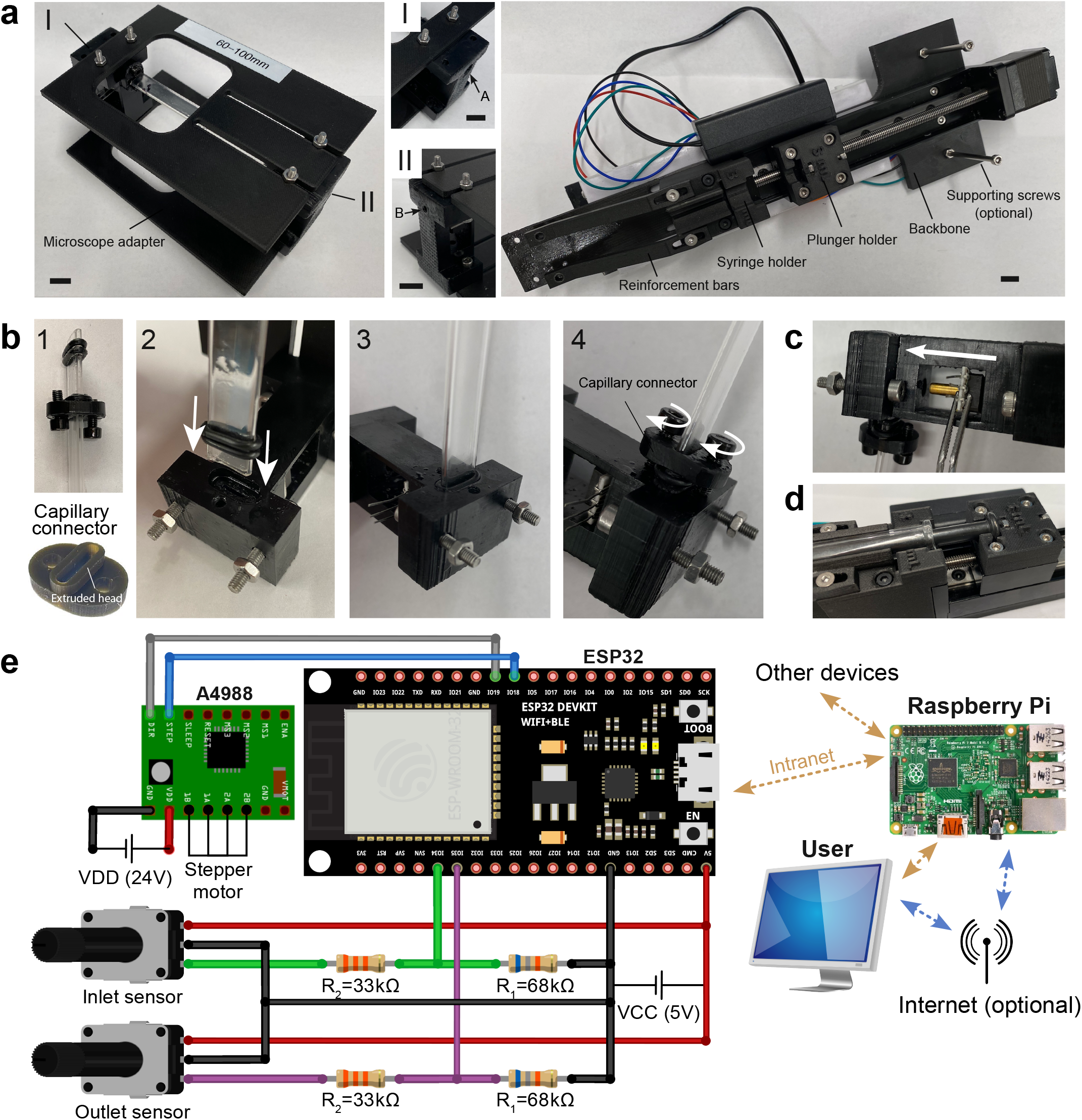
Assembly instructions Installation of the VIEWR. (a) Left: Visualization module with inlet and outlet towers, two rectangular microscope stage adapters (top and bottom) and a glass capillary channel. I: Inlet tower with screw hole A (1/4”-28). II: Outlet tower with screw hole B (M3). Right: syringe pump module with back and assembled linear actuator. Scale bars: 10 mm. (b) Four-step installation of a glass capillary channel to the inlet and outlet towers. (c) Installation of a pressure sensor. (d) Installation of a glass syringe. (e) Wiring diagram. The pinouts shown in this figure are consistent with the attached code, but can be modified if necessary.

The pressure sensors and syringe pump actuators communicate via an embedded ESP32 microcontroller (Espressif Systems) programmed by non-blocking modules (Fig. 2b; more details in Section 2.3). Dataflow is exchanged with the upstream processor (PC or single-board computers, such as Raspberry Pi) with Message Queuing Telemetry Transport (MQTT) protocols through wireless network. A custom Node-RED-based graphical user interface (GUI) is used to facilitate control and data analysis, and has been tested on both laptops (Windows/Mac/Linux) and mobile devices. The dataflow is visualized in real time on the graphical interface and automatically saved locally, which can be readily exported to other readable formats for subsequent processing.

### 2.2 Internet of Things (IoT) architecture

The architecture of our VIEWR system follows the Internet of Things (IoT) paradigm (Fig. 2b). In this architecture, the bottom-level tasks, including actuator control and sensor reading, are packaged into callable functions (written in C++) and executed by the ESP32 microcontroller using its digital and analog pinouts. Users send commands and receive measurement data from a buffer output stored on the microcontroller to enable asynchronous data transfer over wireless communications. This strategic separation of jobs retains actuation and measuring accuracy at an acceptable cost of latency, while further preserving the energy consumed by the microcontroller due to improved workload distribution to minimize intensive computation.

Node-RED, developed and released to open-source communities by IBM in 2016 [26], is a flow-based programming language built based on Node.js and is designed particularly for the implementation of IoT systems. Node-RED provides a visualized programming interface that can be hosted on multiple platforms, including PCs and single microchip units such as the Raspberry Pi. Node-RED is a web-based GUI with an underlying language that is optimized for the easy handling of dataflow, integration of script programming, and functions with a large number of communication protocols that can be embedded into various microcontrollers. Data can be visualized and saved subsequently through wireless communication protocols such as MQTT. Due to the intermediate hosting, users under the same network can access the interface from multiple devices simultaneously, which adds flexibility to the data monitoring and device control.

### 2.3 Non-blocking execution of motor actuation and sensor reading

In a typical rheological measurement, the imposed flow rate and measured analog pressure are controlled and read periodically by the same microcontroller. The flow is actuated by the syringe pump wherein the plunger is mounted to a stepper-motor driven linear actuator. To ensure the accuracy of stepper motor speed, the microcontroller generates alternating digital levels at a precise high frequency. These alternating signals are sent to a stepper motor driver (A4988), in which each rising level triggers a high-voltage impulse to drive the connected motor with one step forward. In contrast, the pressure sensor reading is fed via an analog input and subsequently written to an internal memory buffer for later data transfer, where the latter is more time-consuming. However, the selected ESP32 microcontroller is limited to single-core architecture which does not allow for simultaneous multithreading. As stepper motors only respond to rising digital signals, the microcontroller remains idle at times when the digital output levels are unchanging, and can thus be assigned to perform alternative tasks. This non-blocking scheme to realize multi-tasking is fulfilled by the *BufferedOutput* library (Fig. 2b). Two non-interfering timers and a data buffer in the microcontroller memory are initialized prior to the execution loop. In the background of alternating digital levels (black solid lines, busy in the red shaded area), requests to read the pressure sensors are executed at a user-defined sampling frequency (blue downward arrows), but the results are cached in the internal data buffer and only scheduled to output (blue upward arrows) to the upstream device when the microcontroller reinstates idle (empty areas). Consequently, both operations can be executed simultaneously with minimal interference. Data transmission latency can occur theoretically at the user end due to asynchronous sensor reading and output, but it is negligible in practice, as validated in Fig. 5.

**Figure 4:**
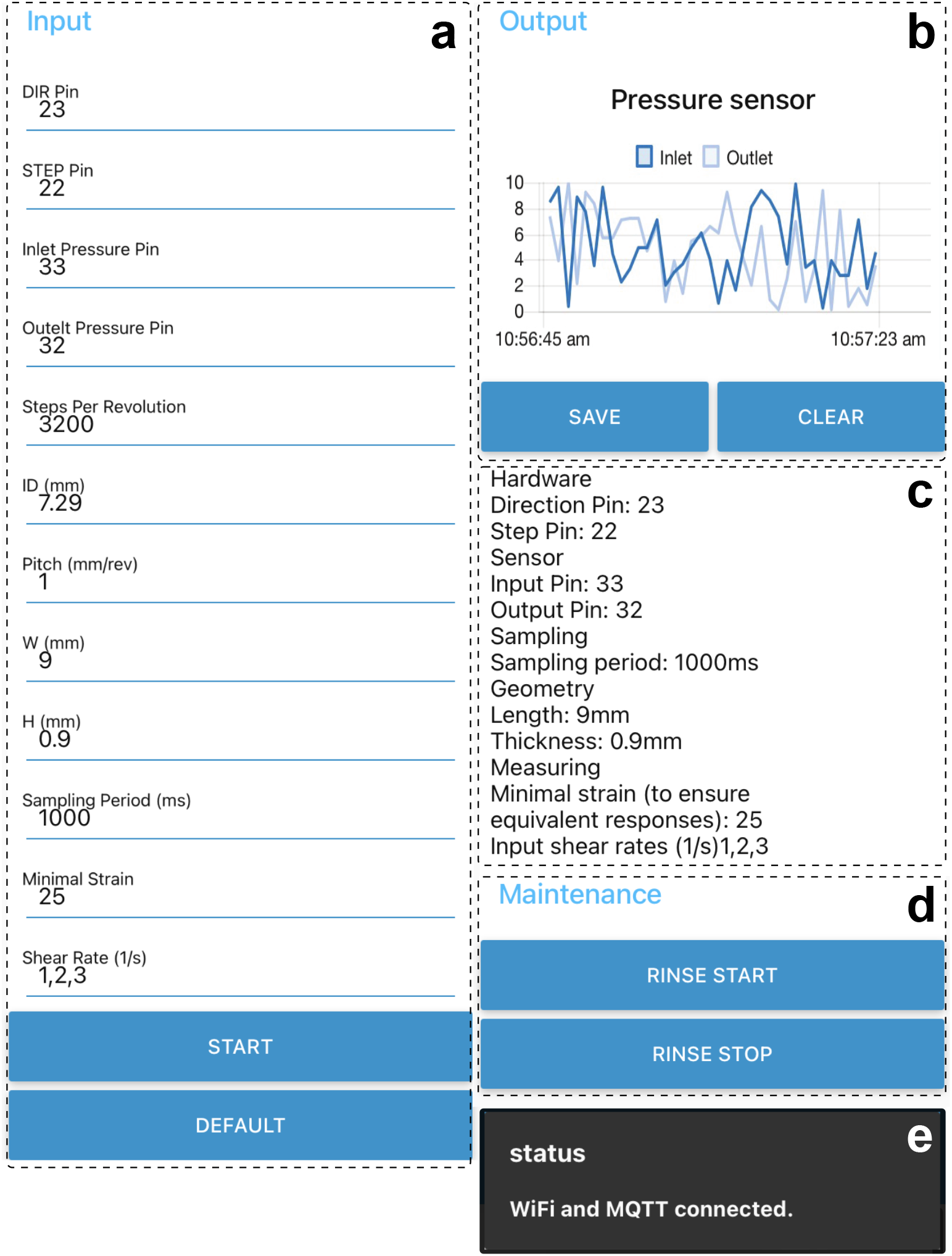
Graphical user interface for VIEWR operation. (a) Input instrument and experimental parameters. (b) Output pressure sensor data for viewing and saving. (c) Overview of current parameters. (d) Maintenance routines for syringe rinsing. (e) User notifications of successful/failed connections or executions.

**Figure 5:**
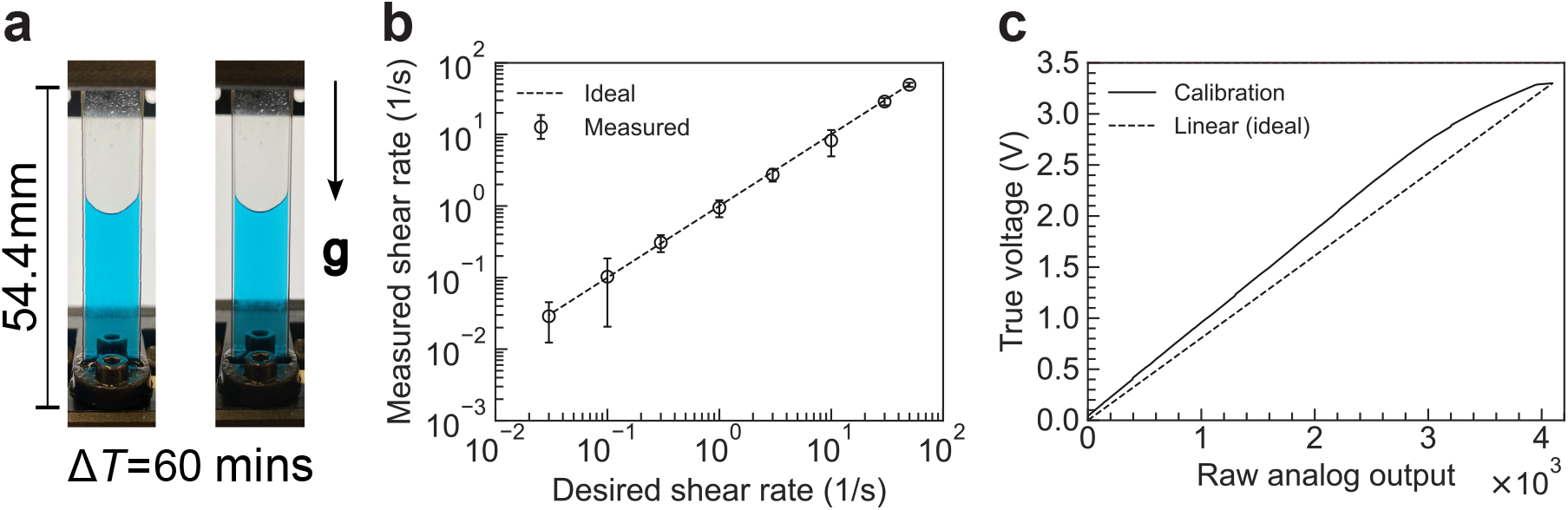
Instrument validation and calibration. (a) Evolution of liquid level in a vertically positioned channel in a duration of 60 mins. (b) Measured shear rates vs. desired values under the non-blocking programming scheme. Error bars correspond to the precision of scale reading on the syringe body. (c) Calibration curve from the raw analog output to the true output voltage. Dashed line: Ideal calibration curve where true voltage is proportional to the analog output.

### 2.4 Key aspects of the design

- VIEWR applies a 3D-printed design for the framework, and uses commercially-available electrical components, resulting in low-cost (approximately $200) manufacturing and easy maintenance.
- VIEWR enables *in situ* visualization of the flow profile and local deformation when mounted on an inverted microscope.
- VIEWR uses interchangeable capillary channels, allowing for measurements with multiple channel sizes to reach a wide range of shear rates. This feature facilitates more stringent pre- and post-experimental cleaning and sterilization, required by certain protocols.
- VIEWR adopts a control/communication protocol through Node-RED and MQTT. This feature enables wireless dataflow and supports experiments in an isolated and well-controlled environment, such as in biosafety cabinets or CO_2_ incubators.

## 3. Design files summary

Design files are summarized in Table 1 with brief descriptions of each file as follows.

- *Framework*: CAD file for the printable framework, which houses all the components.
- *Inlet*: CAD file for the printable inlet tower for the connection of syringe, (inlet) pressure sensor and glass capillary.
- *Outlet*: CAD file for the printable outlet tower for the connection of (outlet) pressure sensor, glass capillary and draining pipe.
- *syringeHolder*: CAD file for the printable syringe holder that connects to the framework and houses the glass syringe for flow control.
- *plungerHolder*: CAD file for the printable plunger holder that connects the slider of the linear rail for flow control.
- *reinforceBar*: CAD file for the printable reinforcement bar that reinforces the connection between the syringe holder and the framework for more accurate flow control.
- *capillaryConnector*: CAD file for the printable connector that joins the inlet and outlet towers with the glass capillary with good sealing and minimal risk of shattering.
- *microscopeAdapter*: CAD file for the printable microscope adapting cover that spans over the inlet and outlet towers to fix on an inverted microscope.
- *Assembly*: CAD assembly file for the guidance of installation and positioning of all individual printable components.
- *motorSensorControl*: Source code (readable by Arduino IDE) preloaded to the ESP32 microcontroller unit for the control of stepper motor and reading of pressure sensors. The Wi-Fi and MQTT credentials in the source code file are left blank and need to be filled in accordingly when adapting to a new instrument.
- *VIEWR:* JSON file to be imported to Node-RED to create the graphical user interface for the VIEWR.

**Table 1:** List of all design files for the manufacturing of VIEWR instrument.

| Design filename | File type | Open source license | Location of the file |
| --- | --- | --- | --- |
| *Framework* | CAD file (.step) | CC BY-NC-SA 4.0 | Framework.step |
| *Inlet* | CAD file (.step) | CC BY-NC-SA 4.0 | Inlet.step |
| *Outlet* | CAD file (.step) | CC BY-NC-SA 4.0 | Outlet.step |
| *syringeHolder* | CAD file (.step) | CC BY-NC-SA 4.0 | syringeHolder.step |
| *plungerHolder* | CAD file (.step) | CC BY-NC-SA 4.0 | plungerHolder.step |
| *reinforceBar* | CAD file (.step) | CC BY-NC-SA 4.0 | reinforceBar.step |
| *capillaryConnector* | CAD file (.step) | CC BY-NC-SA 4.0 | capillaryConnector.step |
| *microscopeAdapter* | CAD file (.step) | CC BY-NC-SA 4.0 | microscopeAdapter.step |
| *Assembly* | CAD assembly file (.sldasm) | CC BY-NC-SA 4.0 | Assembly.sldasm |
| *motorSensorControl* | Source code | CC BY-NC-SA 4.0 | motorSensorControl.ino |
| *VIEWR* | JSON file | CC BY-NC-SA 4.0 | VIEWR.json |

## 4. Bill of materials summary

Bill of materials is summarized in Table 2.

**Table 2:** Bill of materials for the manufacturing of VIEWR.

| Designator | Component | Number | Cost per unit (USD) | Total cost (USD) | Source of materials | Material type |
| --- | --- | --- | --- | --- | --- | --- |
| Printing filaments | Nylon (Onyx™) | 99.8 cm<sup>3</sup> | \$0.24 /cm<sup>3</sup> | \$23.95 | Markforged | Polymer |
| | Isotropic carbon-fiber inlay | 8.64 cm<sup>3</sup> | \$2.90 /cm<sup>3</sup> | \$25.06 | Markforged | Composite |
| SLA resin | B9R-2-Black | 45 cm<sup>3</sup> (est.) | \$0.18 /cm<sup>3</sup> | \$8.10 | B9Creations | Polymer |
| Microcontroller | ESP-WROOM-32 | 1 | \$6.33 /ea | \$6.33 | Link | Other |
| Linear rail | 100 mm | 1 | \$37.44 /ea | \$37.44 | Link | Other |
| Stepper motor controller | A4988 | 1 | \$1.09 /ea | \$1.09 | Link | Other |
| Pressure sensor<sup>a</sup> | BPS130-HG015P-3S | 2 | \$41.81 /ea | \$83.62 | Link | Other |
| Glass capillary<sup>b</sup> | 63832-10 | 1 | \$8.00 /ea | \$8.00 | Link | Other |
| Power components | 9V-battery | 1 | \$2.00 /ea | \$2.00 | Link | Other |
| | 5 V/3.3 V power board | 1 | \$2.00 /ea | \$2.00 | Link | |
| Fasteners and sealers | M3 x 15 (screws and nuts) | 20 | \$0.20 /ea (est.) | \$4.00 | McMaster-Carr | Metal |
| | O-rings (9452K17) | 6 | \$0.03 /ea (est.) | \$0.18 | McMaster-Carr | Plastic |
| | Tube fitting connector (4406T841) | 1 | \$14.41 /ea | \$14.41 | McMaster-Carr | Metal |
| Electronic components | Jump wires | 10 | \$0.10 /ea (est.) | \$1.00 | General supply | Other |
| | Resistors | 4 | \$0.10 /ea (est.) | \$0.40 | | |
| | Breadboard | 1 | \$2.00 /ea (est.) | \$2.00 | | |
| Total: \$219.58 | | | | | | |

<sup>a</sup> Alternative options for pressure sensors can be used if the barb diameter is 3 mm.<sup>b</sup> Alternative sizes of the capillary tubes can be used to reach a wider range of flow rates.

## 5. Build instructions

### 5.1 Hardware

The structural components for the instrument, including the framework, the syringe holder, the plunger holder, the microscope adapter and the reinforcement bars are printed by an extrusion 3D printer (Mark Two, Mark-forged) using nylon filaments, with the addition of two groups of inlayed carbon fiber (three layers each) for mechanical reinforcement. The inlet (I) and outlet (II) towers and the capillary connectors are printed using a stereolithography (SLA) 3D printer (B9 Core 550, B9Creations) which achieves high resolution printing (approximately 100 μm) with biocompatible resins. The SLA printed parts are washed twice in a fresh isopropanol bath for 10 mins each, and are then cured under ultraviolet (UV) light at room temperature for 6 mins to ensure complete polymerization. Prior to assembly, the inlet tower is tapped with 1/4”-28 threading at location A to connect to the glass syringe, and the outlet tower is tapped with M3 threading at location B to connect to the barb adapter (Fig. 3a).

The printed components are assembled according to Fig. 3a-d, followed by the installation of the linear rail. A clean glass capillary is connected to the inlet and outlet towers through the addition of capillary connectors (Fig. 3b). Two capillary connectors are sleeved onto the glass capillary, with the extruded heads facing outwards (Fig. 3b). On each connector, an O-ring is fixed on the extruded head, and another two O-rings are installed directly onto the glass capillary (Step 1). The glass capillary is first inserted into the receiver groove on the inlet and outlet tower until it gently touches the tower surface (Step 2). A tweezer is applied to carefully push the two O-rings on the glass capillary into the gap between the capillary and the receiver groove for proper sealing (Step 3). Finally, the connector is gently pushed into the gap to align the extrusion head with the receiver groove, and the two screws are tightened alternatively with a maximum torque of 0.1Nm to 0.3Nm to avoid shattering the glass. This process is advised to be done by a torque screwdriver. The pressure sensor is installed by first inserting an O-ring into the internal grooves of the inlet and outlet towers (section view of Fig. 2a and Fig. 3c). The sensor head is readily pushed in through the bottom opening without additional fixture. The perpendicularity of the pressure sensor and the flow direction ensures the measurements of purely static pressures. Finally, a syringe filled with deionized water is loaded onto the linear rail for sealing inspection (Fig. 3d), cleaning and optional benchmark measurements, before experiments are performed.

The hardware is connected according to Fig. 3e. All the electrical components can be configured and arranged on a breadboard. VCC can be readily provided by a 9V-battery with a power supply module (listed in Table 2), which also powers the pressure sensors. VDD (24 V) is provided by an external DC power source to drive the stepper motor. Four resistors are used (Fig. 3e) to rescale the output voltages of the pressure sensors from a range of 0-5 V to approximately 0-3.3 V, which is consistent with the readable range of the ESP32 analog pinouts, as well as stabilizing the output voltage, which assists in retaining the measuring accuracy.

### 5.2 Software

The software components of the instrument include a lower-level control for the stepper motor and pressure sensors and an upper-level configuration for data transmission via a wireless communication protocol. The lower-level control is fulfilled by an ESP32 microcontroller unit and an A4988 stepper motor driver. The codefile *motorSensorControl* is attached with this article and must be preloaded to the microcontroller before hardware assembly. To successfully compile the code, *PubSubClient* and *BufferedOutput* libraries are required, which can be installed via the package manager of the Arduino IDE software. In the code configuration, the Wi-Fi information as well as the local IP address for the MQTT host must be filled in prior to the code uploading. The web-based nature of Node-RED allows for a flexible deployment of the network for wireless communications. To optimize the data transmission and security, a Raspberry Pi (or similar portable and linux-based devices) is used as the hosting server (Fig. 3e), which creates an intranet to connect to the ESP32 microcontroller and allows users to control the instrument by directly connecting to the hotspot created by the host server. This architecture allows for the instrument to be operated in the absence of a router or access to the Internet, and can support the connection of multiple microcontrollers or instruments to work collaboratively. The graphical user interface is scripted in JavaScript Object Notation (JSON) and attached with this article. The code can be readily loaded into the Node-RED interface.

## 6. Operation instructions

### 6.1 Connection and graphical user interface

Several preliminary services need to be initialized prior to establishing communications and operating the instrument operation (Table 3). Practically, these steps are configured to automatically launch at server startup, and they remain active while the server is online. Once all services are active, the graphical interface can be readily accessed at `serverhost:1880/ui` from any devices on the same network, where `serverhost` is the IP address of the server that hosts the Node-RED service. The interface consists multiple columns of panels that adapt to the display width (Fig. 4): The first panel (Fig. 4a) is to input the instrument and experimental parameters, which are explained in detail in Table 4. For reviewing purposes, an overview of the input parameters is displayed in a text area (Fig. 4c). A default configuration of the pinouts (according to Fig. 4e) can be loaded by clicking the “DEFAULT” button. During and after measurements, data are streamed and displayed in the display section (Fig. 4b) and can be saved in the format of JSON (“SAVE”) or be cleared (“CLEAR”). Other useful functions, such as rinsing syringes or potential environmental control (temperature or humidity, to be added) can be added in the Maintenance section (Fig. 4d) to facilitate instrument operation in different working scenarios. Finally, a message display space (Fig. 4e) is saved for pop-up notifications of the instrument status, including the connection status of the microcontroller and errors during measurements.

**Table 3:** Prerequisite and initialization of supporting libraries.

| Step | Operation | References |
| --- | --- | --- |
| Wireless communications | An intranet hotspot created by a router or a Debian-based Raspberry Pi using `hostapd` and `dnsmasq`. Wi-Fi credentials need to be consistent with those in the codefile *motorSensorControl*. | [27, 28] |
| MQTT | MQTT service can be turned on by launching `mosquitto` package at port 1883; *Allow_anonymous* must be set **True** to allow for handshaking from other instruments. MQTT IP address needs to be consistent with that in the codefile *motorSensorControl*. | [28, 29] |
| Node-RED | After installing Node-RED, its service is launched by the command `node-red-start`. | [30] |

**Table 4:** Input parameters in the graphical user interface for instrument and experimental configurations.

| Item | Explanation |
| --- | --- |
| DIR Pin | Direction pin for the stepper motor. |
| STEP Pin | STEP pin for the stepper motor. |
| Inlet Pressure | Pin for the analog output of the inlet pressure sensor. |
| Outlet Pressure | Pin for the analog output of the outlet pressure sensor. |
| Steps Per Revolution | Steps per revolution of the stepper motor, which can be controlled by the MS1, MS2 and MS3 pins. |
| ID (mm) | Internal diameter of the syringe. |
| Pitch (mm/rev) | Pitch of the linear rail. |
| W (mm) | Internal width of the glass capillary. |
| H (mm) | Internal height of the glass capillary. |
| Sampling Period (ms) | Time period for each data sampling. |
| Minimal Strain | Minimal strain that each measurement needs to reach to ensure steady-state flow. The time of shearing for arbitrary shear rate $\dot{\gamma}$ equals to (Minimal Strain)/$\max(\dot{\gamma}, 1 \text{ s}^{-1})$. |
| Shear Rate ($\text{s}^{-1}$) | Shear rate(s) for measurements. Multiple shear rates are allowed with the values separated by commas (see example in Fig. 4a). |

### 6.2 Cleaning and sterilization

It is recommended to properly clean and sterilize the channel before taking any measurements to ensure accuracy. To do this, a 15 mL conical centrifuge tube filled with the appropriate and compatible cleaning liquids such as water and 70% ethanol is prepared. A flexible rubber tube (Tube ID is 2 mm) extended from location B in Fig. 3a is inserted into the cleaning liquid, and an empty syringe with the plunger positioned at the minimal volume is installed. By clicking “RINSE START” button on the interface, the cleaning liquid is drawn and extruded alternatively to rinse the channel. The speed and distance of the rinsing can be controlled by the “Minimal Strain” and “Shear rate (1*/*s)” options in the parameter setting column in the interface.

### 6.3 Experimental configuration

After cleaning the instrument, experimental measurements are initialized by loading the fluid sample into the glass syringe, which is subsequently installed onto the instrument. The testing parameters are input in the interface according to the aforementioned instructions. If rheo-optic information is needed, the instrument can be mounted onto an inverted microscope (Fig. 2c and d). The measurements are initialized by clicking the “START” button. During the measuring process, real-time data will be streamed and displayed (Fig. 4b).

### 6.4 Data processing

To minimize the computing workload of the microcontroller unit, only raw analog outputs from the pressure sensors are extracted, and necessary post-processing is required to calculate the material properties.

First, actual pressure values from each sensor can be extracted by a linear calibration relationship as

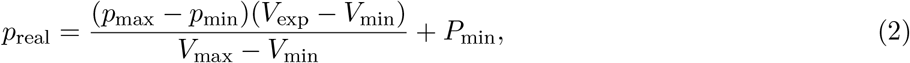

where *p*_max_ and *p*_min_ are the maximal and minimal measurable gauge pressures from the sensors, which are determined in the specifications of the selected sensor. The maximal and minimal voltage outputs, *V*_max_ = 4.5 V are also specified by the sensor manufacturer. The measured voltage *V*_exp_ is extracted from the raw analog outputs (ranging from 0 to 4096) through a calibration function. This calibration is obtained a priori by running a third-party package *esp32-adc-calibrate* using a known base output from a digital pin. More details are presented in the following section, as well as the attached *walkThrough* tutorial (in Jupyter Notebook, Table 1).

Once the pressure measurements from both the inlet and outlet towers are obtained, the material properties and the flow kinematics can be calculated by Eq. 1. However, direct application of this equation does not provide accurate measurements of the shear viscosity. Additional mathematical corrections to the data are necessary to compensate for the finite channel width and the non-Newtonian properties of the tested fluid. Other corrections, such as the entrance effects are shown to have minimal effects on the measuring accuracy within the operational range (Section 7.1), hence are optional but can still be performed to further increase the accuracy.

To incorporate the correction from the finite channel width, the classic analytical solution of a capillary flow through a rectangular channel [31] is applied. Based on the geometry of the applied capillary channel, where *H/W* = 0.1, a simple correction arises as

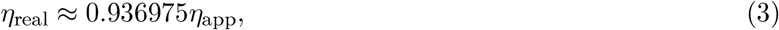

where *η*_app_ is the apparent viscosity calculated based on the 2D simplification, and *η*_real_ is the viscosity that reflects true material properties.

The non-Newtonian effect is corrected by the Rabinowitsch-Weissenberg (R-W) equation [1], where a “true” shear rate for the calculation of material properties can be expressed by the flow rate and the measured pressure difference as

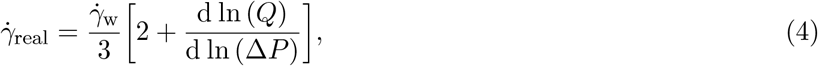

where *Q* is the flow rate. From this equation, for a Newtonian fluid, where Δ*P*∝ *Q*, the “true” shear rate is reduced to the wall shear rate 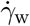.

## 7. Validation and characterization

### 7.1 Instrument validation

The instrument is validated for capillary sealing, the accuracy for the shear rates and voltage calibration for pressure measurements (Fig. 5). To show the proper sealing between the glass capillary channel and the SLA-printed flow tower using the capillary connector (Fig. 3b), the capillary channel is filled with deionized water dyed in blue for better visualization. The height of the water column is monitored, and is found to remain stable over an extended time period of 60 mins (Fig. 5a).

To test the accuracy of flow rates imparted by the syringe pump across a wide range of desired shear rates (from 0.03s^−1^ to 50s^−1^), the stepper motor is driven at a sweeping step frequencies, while both the pressure sensors are sampled at a fixed frequency of 2 Hz. The experimental shear rate is measured by reading the volume change from the syringe body (precision: 20 μL) within a given amount of time. The desired and measured flow rates are in agreement over approximately four orders of magnitude (Fig. 5b), showing negligible artifacts due to the motor stepping or due to interlaced sensor reading during motor actuation.

The ESP32 microcontroller is known to have variations in the reference voltages for analog-to-digital conversion (ADC) [32]. As a result, calibration is needed to convert the raw analog output (read directly by the microcontroller) to the output voltage accurately. Fig. 5c demonstrates the calibration curve from the raw analog output to the output voltage, which deviates from the commonly presumed linear trend (dashed line), especially close to the minimum and maximum limits. Since the raw analog output is discrete, this curve can be readily stored in a look-up table to obtain the voltage values from measured analog outputs without numerical fitting or interpolation.

### 7.2 Measurements on selected material systems

We next sought to validate our VIEWR instrument by measuring the rheology of a variety of materials and comparing our results with those obtained from a commercial oscillatory rheometer (Fig. 6).

**Figure 6:**
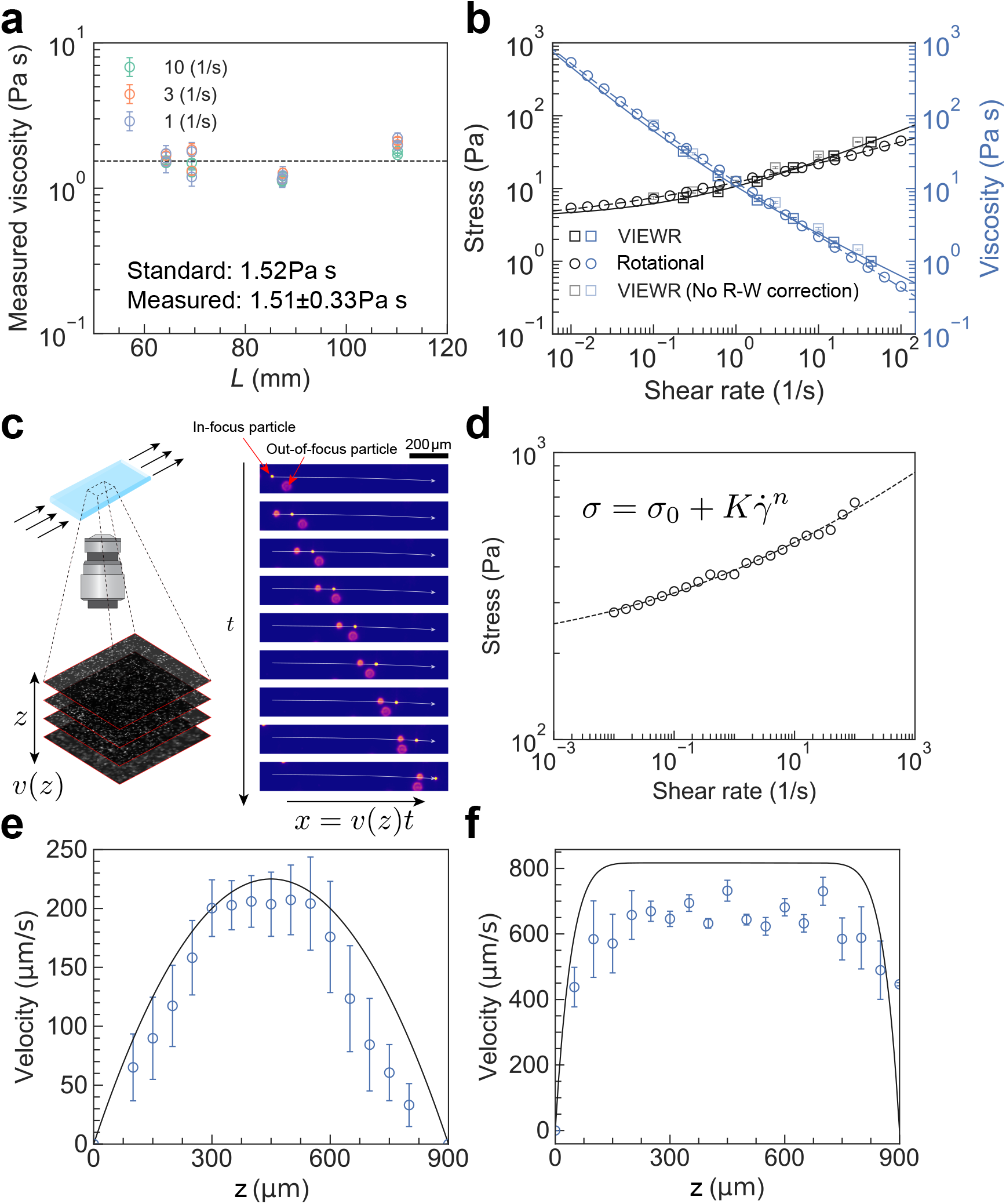
Validation of VIEWR instrument from a selection of materials. (a) Viscosity of silicone calibration oil against the length of capillary channel at varying shear rates. (b) Flow curves of aqueous carbopol solution (0.1w%) measured from VIEWR (square markers) and rotational shear rheometer (circle markers). The shaded square markers represent measurements without Robinowitsch-Weissenberg (R-W) corrections for reference. Solid and dashed lines represent fitting lines to the Herschel-Bulkley model of the measurements from VIEWR after R-W corrections and the rotational rheometry, respectively. (c) Schematic representation of the *in situ* microscopy to determine the flow profile, where image processing algorithms are used to determine the in-focus features for particle image velocimetry (PIV). (d) Shear rheology of aqueous pluronic solution (27w%) at 25 °C measured from rotational rheometer. Dashed line represents fitting line to the Herschel-Bulkley model. (e) Theoretical (black line) and measured (blue markers) velocity profiles of glycerol at an imposed wall shear rate of 1s^−^. Errorbars represent the standard deviations of velocity measurements from legitimate particle trajectories (with a number of 5 to 10 depending on the focal plane). (f) Theoretical (black line) and measured (blue markers) velocity profiles of aqueous pluronic solution (27w%) at 25 °C at an imposed wall shear rate of 5 s^−1^.

First, benchmark tests were performed with a Newtonian viscosity standard (S600, Cannon Instrument) at 23¶C (Fig. 6a). We measured its steady-state viscosity at various flow rates of 10s^−1^, 3s^−1^ and 1s^−1^, incorporating the geometry correction (Eq. 3). To quantify the entrance/exit effects from capillary rheometry, we performed additional measurements with varied capillary lengths ranging from 65 mm to 110 mm. The mean value of the measured viscosity ((1.51 ± 0.33) Pas) are in excellent agreement with the standard value of 1.52 Pa s (black dashed line) without significant deviation. As a result, the entrance/exit effects from the VIEWR instrument can be ignored when the material systems are viscous (≥1Pas).

To demonstrate accurate measurements of a non-Newtonian fluid from the VIEWR instrument, especially bioinks or material with similar rheological properties, a yield-stress fluid (0.1w% Carbopol 1040 in DI water neutralized with NaOH, Lubrizol Corporation) was tested over a shear-rate range of 0.1 s^−1^ to 30 s^−1^. Carbopol has extensive applications in biology, pharmaceutical development and embedded 3D bioprinting [33, 34], and rapid measurements of the yield-stress are of great importance to understand and optimize for material behaviors. As shown in Fig. 6b, the measurements of shear stress and viscosity before (shaded squares) and after (dark squares) Robinowitsch-Weissenberg corrections are plotted against the shear rates. The two results are further compared with independent rheological measurements (circle markers) on a rotational rheometer (ARES-G2, TA Instruments). We extracted the yield stress by fitting the measurements from the capillary rheometry (after R-W corrections, solid lines) and the measurements from a rotational rheometer (dashed lines) into the Herschel-Bulkley model, from which the stress can be expressed as

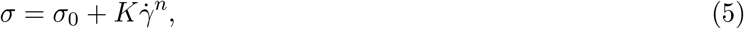

where *σ*_0_ is the yield stress, and *K* and *n* describe the power-law shear-thinning trend at large shear rates. The fitted results show *σ*_0_ = 3.9Pa from the corrected measurements, and *σ*_0_ = 3.2 Pa from the rotational rheometry, which are in excellent agreement.

To measure the velocity profiles of complex fluids through direct observation, we tested two material systems which find extensive applications in 3D printing: glycerol (Fig. 6e) and pluronic solutions (27w%) (Fig. 6f). To facilitate the particle image velocimetry (PIV), 10-μm fluorescent particles (FluoSpheres F8836, ThermoFisher Scientific) were added to the tested samples at an overall particle density of 3.6 × 10^4^ mL^−1^ and homogenized with a speedmixer (FlackTek Manufacturing Inc.) at 1500 rpm for 15 s. To measure the flow profile, the VIEWR instrument is installed on an inverted microscope with a 4x objective (MRL00042, Nikon Instruments Inc.), and a constant shear flow is initiated. The objective is focused on the bottom plane of the internal channel and a z-stack is acquired with 50 μm spacing. At each slice, a series of snapshots are captured by a global-shuttered sCMOS camera (Kinetix, Teledyne Photometrics) at a preset frequency, with synchronized 488-nm excitation laser pulses (Fig. 6c).

The resulting image series are post-processed by a third-party imaging processing package *trackpy* [35]. To distinguish the velocity profile at specific focal planes, only in-focus particles filtered based on their size, mass, and peak brightness are used for velocimetry.

For glycerol, the theoretical 2D flow profile (solid line, Fig. 6e) is expected to be parabolic and is solely a function of the imposed flow rate but independent of the material properties. In contrast, the aqueous pluronic solution is characterized as a yield-stress fluid (Fig. 6d) from independent rotational rheometry, and its flow behavior can be well fitted by the Herschel-Bulkley model (dashed line) with the yield stress *σ*_0_ = 197.2Pa. The flow profile predicted from the Herschel-Bulkley model based on their constitutive parameters is calculated from Cauchy’s momentum equation and the boundary conditions, and exhibits a highly non-parabolic profile with plug-flow features (no velocity gradient) near the center (solid line, Fig. 6f). For both sample fluids, the extracted flow profiles (blue markers) are in good agreement with the predictions, despite lower magnitude in the peak velocity from the measurements, which may arise from the image processing that failed to distinguish out-of-focus particles. Overall, the consistency between the flow-profile predictions and measurements demonstrate satisfying accuracy for flow control and *in situ* characterization by the VIEWR instrument.

## Ethics statements

No experiments on human or animal subjects were performed in this article.

## CRediT author statement

**Jianyi Du**: Conceptualization, Design, Manufacturing, Measurement, Visualization, Data processing, Writing. **Soham Sinha**: Design, Hardware, Manufacturing. **Stacey Lee**: Design, Measurement, Writing. **Mark A. Skylar-Scott**: Conceptualization, Writing, Supervision.

## Acknowledgements

J.D. and M.A.S.-S. thank the financial support from Stanford Children’s Heart Center. This research was funded in part by a Chan Zuckerberg Investigator Fellowship. S.S. is supported by the National Science Foundation Graduate Research Fellowships Program (Award Number DGE-1656518) and Benchmark Stanford Graduate Fellowship. J.D. and M.A.S.-S. thank the technical help on the high-speed imaging system from Prof. Polly Fordyce at Stanford University, as well as the insightful discussions with Caroline Horn, Peter Suzuki and Michael Hayes on the experimental design and analysis. Part of this work was performed at the Stanford Nano Shared Facilities (SNSF), supported by the National Science Foundation under award ECCS-2026822.

